# Controlling Cellular Arrangements via Stretched Bioprinting

**DOI:** 10.1101/2020.11.30.403378

**Authors:** Chuanjiang He, Mengxue Liu, Deming Jiang, Chunlian Qin, Tao Liang, Pan Wu, Chunmao Han, Liquan Huang, K. Jimmy Hsia, Ping Wang

## Abstract

Bioprinting is a common method to replicate geometrical architecture of native tissues. However, it usually fails to modulate cellular arrangements, which is critical for the tissue’s functionality. To our knowledge, no method has successfully addressed this challenge. Here, we report a method of controlling cellular orientation during the bioprinting process by integrating a stretch process into a modified bioprinting frame. We demonstrate that the cellular orientation is a result of cells’ sensing and responding to the tensile stress, instead of shear stress or topographical patterns. Moreover, our method shows a potent capability to induce myoblast differentiation, fusion and maturation without the presence of differentiation medium. As a potential clinical application, we demonstrate that aligned myofibers directly printed onto injured muscle in vivo, can not only repair the structure of damaged tissue, but also recover the muscle functionalities effectively. This study shows that the new method can produce tissues with precise control of cellular arrangements and more clinically viable functionalities.

**Significance Statement:** Due to no method could reproduce the exact cellular arrangements of native tissues in engineered tissues, tissue engineering facing difficult in fabricating 3D tissues that possess desirable biological and mechanical functionalities for biomedical applications. For the first time, we report a method of controlling cellular orientation during 3D bio-printing process. This method can be used to produce engineered tissues with controlled cellular arrangement with several different cell types. Moreover, this method shows a potent capability of fabricating fully mature and aligned myofibers *in vitro* in the absence of differentiation medium. As potential clinical applications, with this method, engineered tissues could be directly printed *in vivo* with high efficacy of tissue repair and function recovery.

## Introduction

3D bioprinting has shown potentials in producing viable tissue replicates or artificial organs, such as heart (1), vascular grafts (2), multilayered skin and tracheal splints (3), which can be used for transplantation, drug discovery and in vitro disease study (4). Although these cellular constructs possess similar shape and dimension as the real tissues or organs, they usually fail to reproduce the cellular arrangements of native tissues (5). For cells and tissues in their native environment, such as neurons, tendon, vascular tissues or striated muscle tissues, their biological and mechanical functionalities are dictated by the intricate cellular arrangements, e.g., cell alignment (6, 7). For instance, the arrangements of myocardial cells and skeletal muscle cells are critically important for cardiac beats and muscle contraction, respectively (7, 8). Thus, it is of great importance to control the orientations of cells in the manufactured tissues, and simple improvements as controlling cellular alignment in engineered tissues would help recapitulate the functionalities of damaged tissues.

There is growing recognition that cellular responses to biophysical stimuli, such as topographical patterns or mechanical strain play a crucial role in cellular orientation/re-orientation, although the mechanism is not fully understood (6, 9). It has been shown that adherent cells can sense and align with micro- or nanoscale anisotropic patterns of the substrate (6, 10, 11). Fabricating topographic structures in bio-ink for printing, however, is impractical. Cyclic or static stretching of engineered tissues can also induce the alignment of cells, as demonstrated by clamping shape-specific tissues in mechanical stretch bioreactors (12). This technique, however, is only suitable for engineered tissues with certain strength and specific shape and size. Moreover, the stretching direction of the bioreactors is limited to two fixed directions, thereby insufficient in manufacturing physiologically relevant structures

In the current study, we have developed an effective bioprinting method to control the cellular orientation while fabricating cellular constructs. We re-arranged the extrusion bioprinting process from conventional “extrusion-deposition-gelation” to “gelation-extrusion-stretch-deposition”, and thus controlled stretching strain could be applied to bioink (**Fig. 1A**). With a simple customized handheld device, we demonstrate printing of patterned 3D constructs of various cell types (cardiomyocytes, cardiac fibroblasts, airway smooth muscle cells (ASM), human skin fibroblasts (HFF-1), dorsal root ganglion (DRG) neurons and myoblasts (C2C12)), in which the cells align either in the interior or on the surface of the hydrogel. Furthermore, we show that the cellular alignment is the result of cells sensing and responding to the state of tensile stress, instead of shear stress or topographical patterns on the substrate. Moreover, the results indicate that cells prefer to divide and elongate along the direction of the maximum tensile stress. Using this technique, we are able to construct, *in vitro,* nearly 100% mature and perfectly aligned engineered myofibers, which form in the absence of differentiation medium to induce myogenesis. We also demonstrated, *in vivo*, the ability of the printed aligned myofibers to repair injured muscles and recover the structure and functionalities of volumetric muscle loss.

**Fig. 1.**
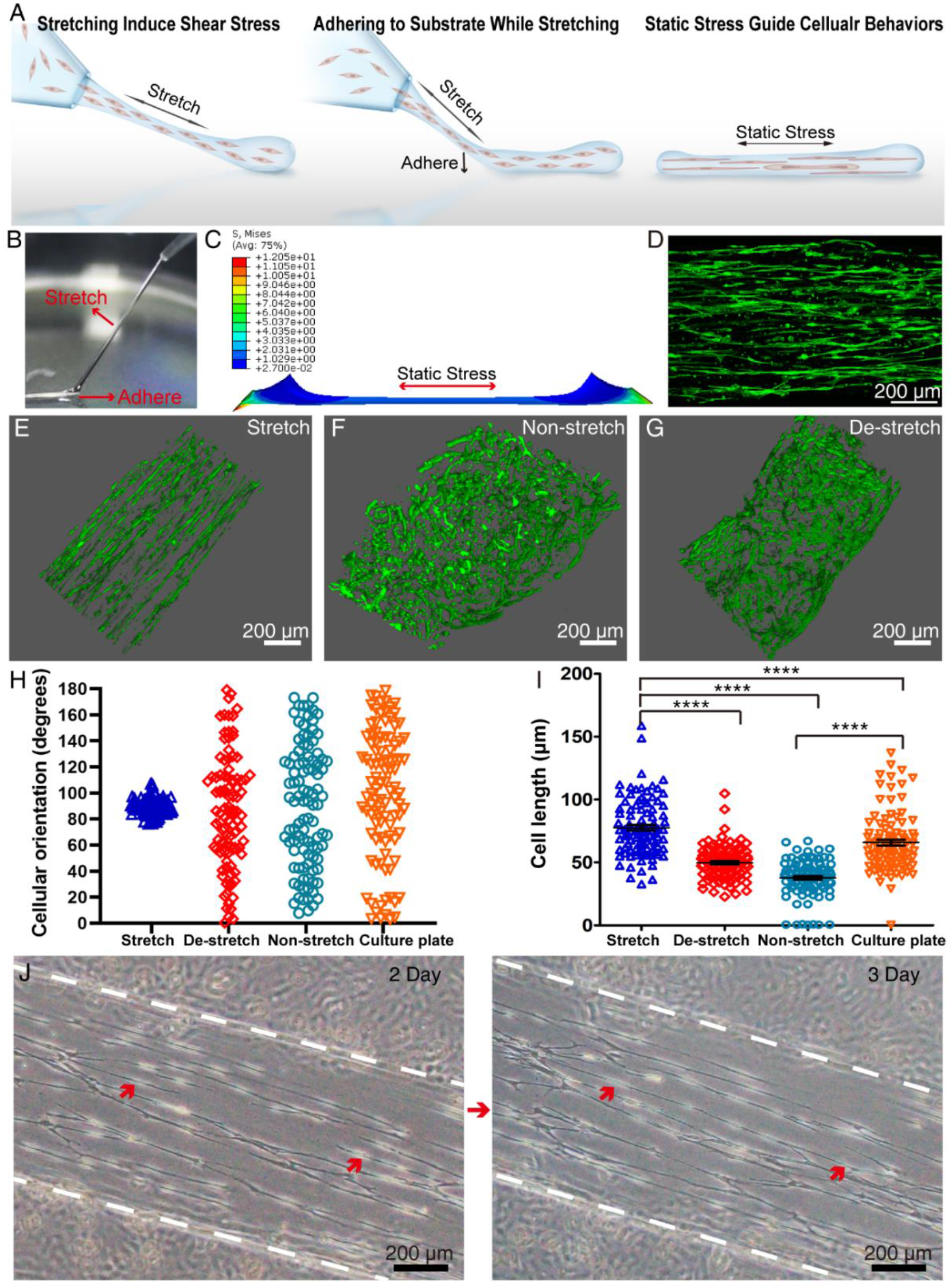
Controlling cellular orientation via stretch-bioprinting. Schematic (**A**) and photograph (**B**) depicting the printing process. (**C**) Simulation result of the static stress distribution in the printed gelatin filament. (**D**) Fluorescence image of cells aligning in the generated constructs. Confocal fluorescence 3D images of cellular arrangement generated by printing with stretch (**E**), non-stretch (**F**) and de-stretch (**G**) process. The distribution of cellular orientation angles (**H**) and cell length (**I**) in 3D constructs generated by different printing process. The 90 angle represents the direction of stretching and the culture plate group represents cells seeded onto 6-well culture plate. One-way ANOVA with Tukey’s multiple comparisons test, n=100, Error bars: means ± SEM, ****p < 0.0001. (**J**) Dynamic cell division process of cardiac fibroblasts that were seeded onto the surface of stretched gelatin, and the newborn cells bridged the gap between arrows on the third day. Green: phalloidin- iFluor 488.

## Results

### Stresses induced by stretching are critical for controlling cellular orientation

In this study, we re-arranged the extrusion bioprinting process from conventional “extrusion-deposition-gelation” to “gelation-extrusion-stretch-deposition”. In detail, prior to the extrusion step, gelation of hydrogel-cell mixture was induced in a microtube to form uniform cylindrical gel filament. The gelation of hydrogel started about 12 minutes after mixing gelatin with microbial transglutaminase (mTG) and cells. After about 25 minutes, the gel became sufficiently stretchable to sustain large strain during the bioprinting process (Fig. S1). Then, gel filament (about 650-750 μm in diameter) was extruded from the microextrusion head. While the end of the gel filament was adhered to the substrate, the gel filament was stretched and fixed by horizontal and vertical moving the microextrusion head relative to the substrate, respectively (Fig. 1B). When adhered to the substrate, the stretching strain in the gel filament was maintained. The finite element analysis (FEM) simulation in Figure 1C shows that the tensile stress in the adhered gel filament is uniform along the filament.

In experiments with cardiomyocytes, stretching strain levels from 100% to 400% (Fig. S2) were applied to the printed gelatin filaments. The results showed that nearly all cells aligned in the direction of stretching in the interior or on the surface of the 3D gel without impairing their contractility (Fig. 1D, Movie S1), demonstrating the technique’s ability to generate well-aligned, 3D cellular constructs. The elongation of the cells was weakly dependent on the stretching strain.

Key questions here are what role the post-extrusion stretching plays in cell alignment, and what the driving force underlying such cell arrangement process is. To answer these questions, we performed the control experiment of extruding the gel filament without stretching. We also attached the stretched filaments to a PDMS substrate under “dry” condition where the filament tended to detach from the substrate when the culture medium was added after about 20 minutes. Once detached, the stretching stress is relaxed, which we called the “de-stretched” condition. Cell responses under these conditions were examined. Fig. 1E clearly shows that cells completely aligned themselves in the presence of stretching. In the control experiment of the extruded filament without stretching, the cardiomyocytes did not show alignment as shown in Fig. 1F. Furthermore, under de-stretched condition, the cardiomyocytes again exhibited random orientations (Fig. 1G). The results under de-stretch condition suggest that the persistent presence of tensile stress is essential for inducing cellular alignment. Under persistent presence of the static stretching, cells always elongate and align in the axial direction of the filament, i.e., the maximum tensile stress direction (Fig. 1H, I). Moreover, time lapse observations showed that the dynamic cell division also tends to occur along the maximum tensile stress direction (Fig. 1J), consistent with the results of other studies (13–16).

### Role of shear stress and anisotropic topography in cellular arrangement

The hydrogel embedded with cells is a viscoelastic material, undergoing not only elastic but viscous deformation during stretching. In particular, the stretching may reduce the local viscosity of hydrogel due to fast dissociation of the crosslinked complexes, a phenomenon referred to as shear-thinning (17). During stretching, the hydrogel undergoes microscopic reconstruction of crosslinked networks. Therefore, there is a transition of gel-dynamic solution-gel in hydrogel. This microscopic reconstruction gives rise to an effective shear stress in the hydrogel filament, but the shear stress disappears during steady state stretching. During this transient phase, the shear stress may drive re-alignment of anisotropic objects, such as cellulose nanocrystals (17) (Fig. 2A). We investigated it by stretch-printing gelatin embedded with silver nanowires, and parallel alignment of the silver nanowires was observed (Fig. 2B). Such effect is not only observed in entities with large aspects ratio such as fibers, but also noticeable in non-elongated cells. However, Fig. 2C, taken shortly after the stretching, shows that such shear stress effect cannot account for the cell alignment observed upon long-term stretching. To further investigate this, we seeded cells onto the surface of stretch-printed gelatin to eliminate the influence of shear stress. 45 minutes after being seeded, cellular pseudopodia started to form along the direction of stretching (Fig. 2D), indicating that cells could sense the direction of stretching without the presence of shear stress. After 2 days of culture, the cells seeded onto the surface of printed gelatin filaments were still aligned and were more elongated than those seeded onto the substrate of culture plate (Fig. 2E). In contrast, when seeded onto the surface of a de-stretched gelatin filament, cells show no alignment and less elongation (Fig. 2F).

**Fig. 2.**
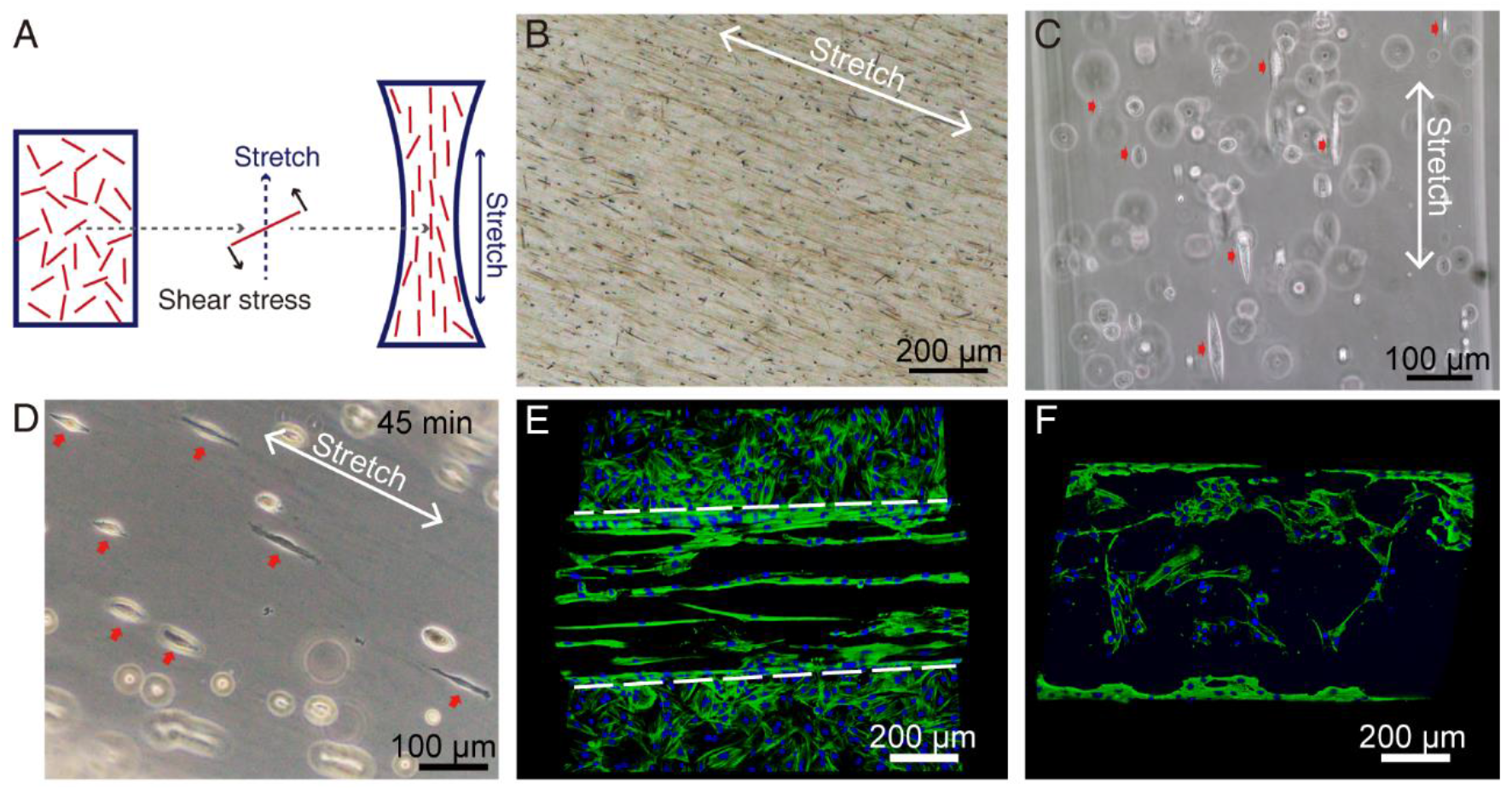
Role of shear stress in cellular arrangement. (**A**) Schematic illustration of the hydrodynamic alignment of anisotropic substance induced by shear stress. Images of silver nanowires (**B**) and unspreading cardiac fibroblasts (**C**) oriented longitudinally in the gel induced by shear stress. (**D**) Cardiac fibroblasts were seeded onto the surface of stretched gelatin filament for 45 min and formed pseudopodia along the direction of stretching. Fluorescence images of cardiac fibroblasts that were seeded onto stretched (**E**) and destretched (**F**) gelatin filaments for 2 days. Green: phalloidin- iFluor 488; Blue: DAPI.

Considering that cells can sense and align with anisotropic topography (6), we wondered whether contact guidance plays a role in our system. However, no anisotropic patterns were observed in the inner and on the surface of lyophilized specimen of stretched gelatin (Fig. S3A, S10B). Furthermore, with the mechanism that air-drying of stretched gelatin strips could cause further lateral volume contraction and surface instability (17, 18), we fabricated surface wrinkles onto the stretched gelatin strips (Fig. S3C). Then, we seeded cells onto stretched gelatin with and without wrinkles, and cells preferred to fall into grooves after seeded (Fig. S3D). However, as to cellular orientation, there are not any difference between them (Fig. S3E, S3F). These results suggested that there was no impact of contact guidance in our system.

These observations suggest that cellular orientation and elongation in this system are the direct result of cells responding to the persistent presence of tensile stress that generated by stretching. Because the cytoskeletal proteins such as actin and tubulin are crucial for mediating cellular responses to mechanical cues (9), we try to block these responses by using cytoskeleton-disturbing chemicals, cytochalasin B and blebbistatin, to disrupt cellular filamentous actin. As a result, the number of cells adhering and aligning onto the stretched gelatin dropped significantly, since the adhesion and spread of cells were inhibited (Fig.S4).

### Cell alignment in patterned constructs

One of the biggest challenges in tissue engineering is being able to generate functional tissues that conform to the complex 3D shape of organs in the human body. Although there are techniques to produce live cell constructs of conformal shapes such as scaffolding, the cell arrangement in these constructs cannot be controlled, leaving the engineered tissues less functional. In this study, we demonstrate that the current method can tissues of complex shapes with aligned cells. As shown in Fig. 3, several 3D letter-shaped cell constructs were printed with cells aligned following the longitudinal direction of the filament.

**Fig. 3.**
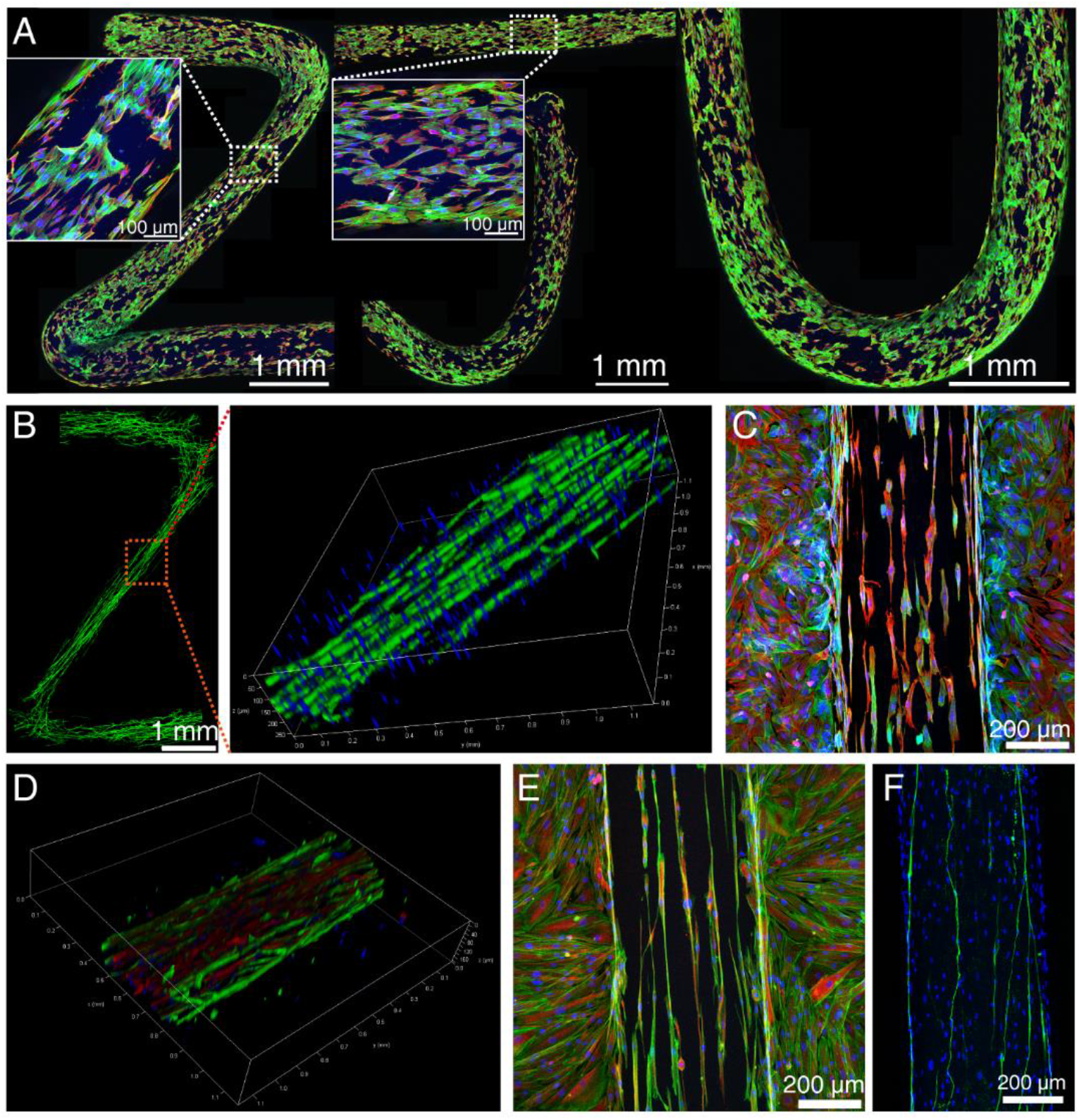
3D patterned constructs with several types of cells aligned in the interior or on the surface. (**A**) Maximum z-projection image of cardiac fibroblast [marked with antibody to vimentin (red)] that aligned onto the surface of printed 3D letter-shaped gelatin constructs. (**B**) Confocal fluorescence 3D images of ASM cells aligning in the Z-shaped gelatin construct. (**C**) Maximum z-projection image of ASM cells [marked with antibody to alpha smooth muscle actin (red)] that were seeded onto the surface of stretched gelatin for 2days. (**D**) Confocal fluorescence 3D image of HFF-1 [marked with antibody to vimentin (red)] aligning in the inner of stretched gelatin. (**E**) Maximum z-projection image of HFF-1 cells [marked with antibody to vimentin (red)] that were seeded on the surface of stretched gelatin. (**F**) Rat dorsal root ganglion neurons [marked with antibody to β-III tubulin (green)] aligned orderly along the stretched gelatin. Green: phalloidin- iFluor 488; Blue: DAPI.

Our method can induce cell alignment not only in cardiomyocytes, but also in several other cell types. Figures 3A-F, S5, and Movie S1 show the alignment of several cell types in gelatin filaments, including rat cardiomyocytes (Fig. S5, Movie S1), rat cardiac fibroblasts (Fig. 3A), rat airway smooth muscle cells (ASM) (Fig. 3B, 3C), human skin fibroblasts (HFF-1) (Fig. 3D, 3E), and rat dorsal root ganglion neurons (Fig. 3F). Furthermore, depending on the adhesive property of the stretched gelatin, individual gelatin filaments with different orientations may form adhesive junctions with each other. In such cases, more complex, 3D stacked constructs can be produced by several filaments with the cellular orientations controlled by the 3D printing process (Fig. S6). Moreover, photo-crosslinked gel, such as gelatin methacrylate (GelMA) also can be used as bioink in this method (Fig. S7).

### Fully mature and aligned myofibers formation in the absence of differentiation medium

In the skeletal muscles, normal myofiber tracts are composed of highly ordered and aligned bundles of multinucleated myotubes formed by the fusion of myoblasts (19) (Fig.4A). In *in vitro* culture, myotube formation requires differentiation medium over certain duration. If the muscle cells are initially randomly oriented, the resulting myotubes would also be oriented randomly. Using our method, however, we were able to produce aligned and mature myofibers with myomimetic hierarchical structure *in vitro*, and without adding differentiation medium, as shown in Fig. 4B and Movie S2. With our method, fully mature mytotubes formed on Day 8 without further manipulation.

**Fig. 4.**
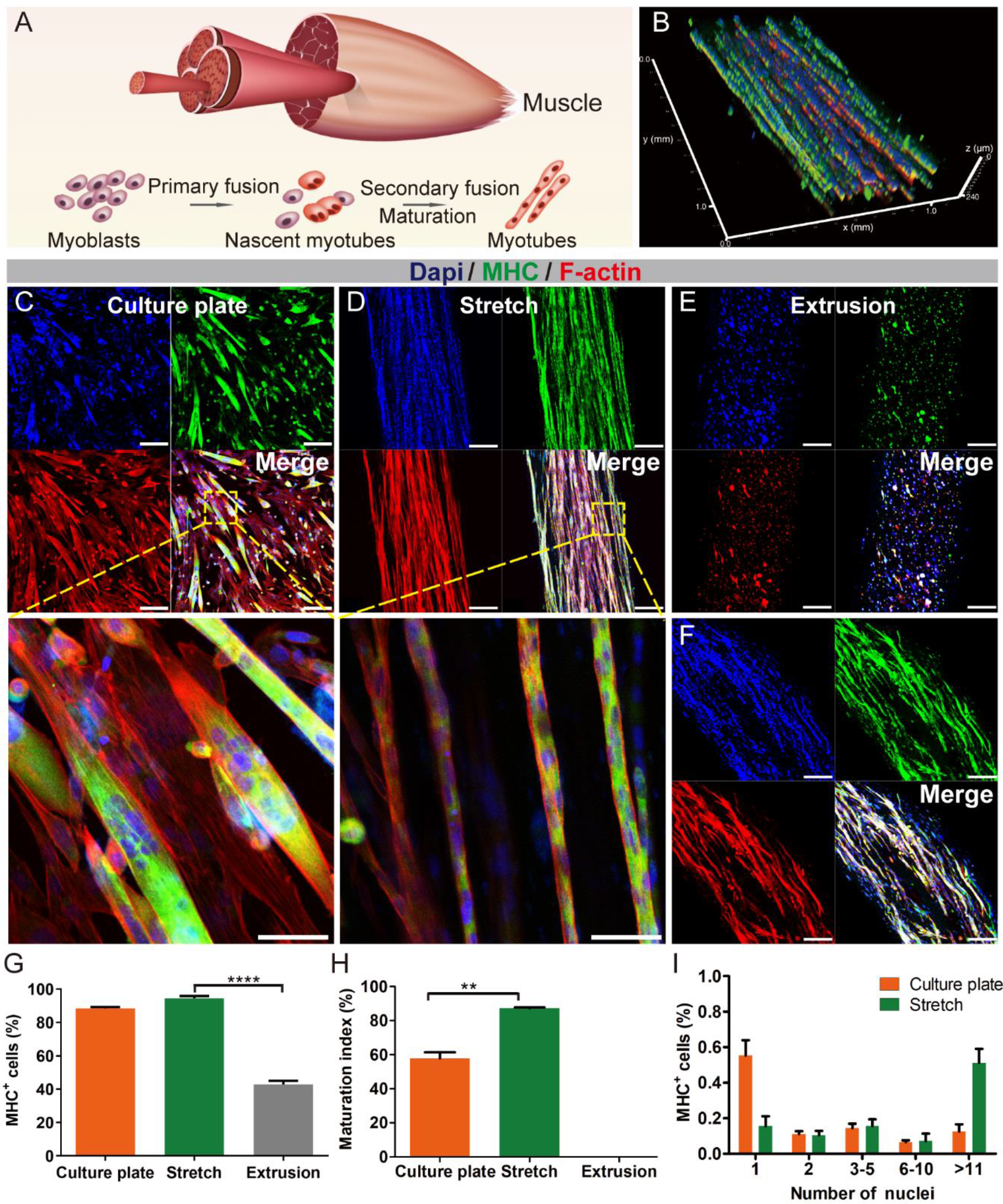
Fabricate aligned mature myofibers in vitro. (**A**) Schematic illustration of the structure of muscle and the formation process of myotubes. (**B**) Confocal fluorescence 3D images of the myomimetic structure of printed myofibers. (**C**) Confocal fluorescence images of C2C12 cells that were seeded onto culture plate for 8 days (added medium to differentiation medium on Day 4), bar= 200 μm. Below: magnified image, bar= 50 μm. (**D**) Maximum z-projection images of aligned myotubes in the printed gelatin (without using differentiation medium), bar= 200 μm. Below: magnified image, bar= 50 μm. (**E**) Maximum z-projection images of C2C12 cells in the as-extruded gelatin without stretching. Bar= 200 μm. (**F**) Maximum z-projection images of C2C12 cells in the gelatin upon destretching on Day 8. Bar= 200 μm. MHC positive index (**G**) (one-way ANOVA with Tukey’s multiple comparisons test, ****p < 0.0001), maturation index (**H**) (unequal variance Student’s t-test, **p =0.0012) and the distribution of the number of nuclei in MHC positive cells (**I**). Green: antibody to myosin skeletal heavy chain (MHC); Red: phalloidin-YF555; Blue: DAPI. Error bars: means ± SEM.

In this study, the behaviors of C2C12 mouse myoblasts were examined. C2C12 myoblasts differentiate into myotubes upon adding differentiation medium when cultured in a petri dish as shown in Fig. 4C., Interestingly, Fig. 4D shows that C2C12 cells in stretched gelatin differentiated, without differentiation medium, into perfectly aligned myotubes with greater fusion and maturation than those in the petri dish, and without the damnification of cell proliferation (Fig. S8). These axially aligned engineered microfiber bundles exhibit phenotype very similar to that of native muscle fibers. This observation raises an important question about how, in addition to biochemical cues such as those in differentiation medium, mechanical environment can directly influence or induce myogenic differentiation, fusion and maturation. We also did the control experiment on as extruded filament without stretching. Figure 4E clearly shows that, in the as-extruded filament without stretching and without differentiation medium, few (if any) cells showed tendency of fusion, maturation, and myotube formation.

We further observed that, when we detached the printed gelatin from substrate on the 7th day, the myofibers exhibited elastic spring-back and curvilinear shapes (Fig. 4F). This suggests that, when the filaments adhering to the substrate, the myofibers were in a tensile stress state, i.e., a tensile stress is essential for myotube formation. Quantitative measurements of the myosin skeletal heavy chain (MHC) positive index (the ratio of the number of positive nuclei in the MHC to the total number of nuclei) shown in Fig. 4G, the maturation index (the ratio of the number of myotubes consisting of more than five nuclei to the total number of myotubes) in Fig. 4H, and the distributions of the serial numbers of nuclei in MHC positive cells in Fig. 4I all indicate that our method can more efficiently induce myoblasts fusion and maturation (nearly 100%) compared with conventional 2D culturing plus differentiation medium. These results showed the power of mechanical cues at inducing myofiber formation in vitro.

### Application: aligned myofibers to repair damaged muscle *in vivo*

It is important to know whether the gelatin filaments with well-aligned myofibers produced by our method would be a better artificial tissue for wound healing or damaged tissue repair. We explored this issue by directly printing stretched gelatin filaments loaded with C2C12 myoblasts onto damaged mice tibialis anterior muscles. Using a volumetric muscle loss (VML) model, we removed about 30% volume on a tibialis anterior muscle tissue (Table S1). We then stretch-printed C2C12-loaded gelatin filaments on the wound directly, and aligned filaments with the direction of the native muscle fibers (Fig. 5A, 5B and Movie S3, Trypan Blue was added to enhance the visibility of gelatin). Four controlled experimental groups were performed (n=5): (1) untreated wound; (2) stretched gelatin without embedded C2C12 myoblasts (Gel+Stretch); (3) gelatin scaffolds loaded with C2C12 myoblasts without stretching (Gel+Cells); (4) stretched gelatin filaments with C2C12 myoblasts aligned with the direction of muscle fibers (Gel+Cells+Stretch). Evidence showed that the functional performance of mice treated with stretched gelatin with C2C12 (Group 4) was better than that of the other groups in four limb hanging tests and rotarod running tests (Fig. 5C, 5D). Moreover, injured muscle by Group 4 treatment exhibited significant growth of muscle mass (Table. S1), while other three control groups contained significant fibrotic tissue and reduced myofibers at the wound sites (Fig. 6A, 6B). It was also observed that, for the wound treated with stretched gelatin without cells or as extruded cells-loaded scaffolds (Groups 2 and 3), there was still unabsorbed gelatin after 3 weeks (Fig. 6A, 6B). In contrast, for wound treated with Group 4 filament, newly regenerated myofibers (centrally nucleated MHC+ myofibers) and fewer fibrotic tissue were evident in Figs. 6B and 6C, and the gelatin was completely absorbed (Fig. 6A, 6B). These regenerated myofibers exhibited organized circular laminin structure similar to that in normal muscles. In comparison, wound treated by Groups 2 and 3 showed disorganized laminin deposition within the wound sites (Fig. 5D). Furthermore, the wound area treated by Group 4 was populated with more robust and denser capillary networks than the other groups (Fig. 5E). These results clearly demonstrate that our method could present a powerful technique for tissue regeneration and tissue repair.

**Fig. 5.**
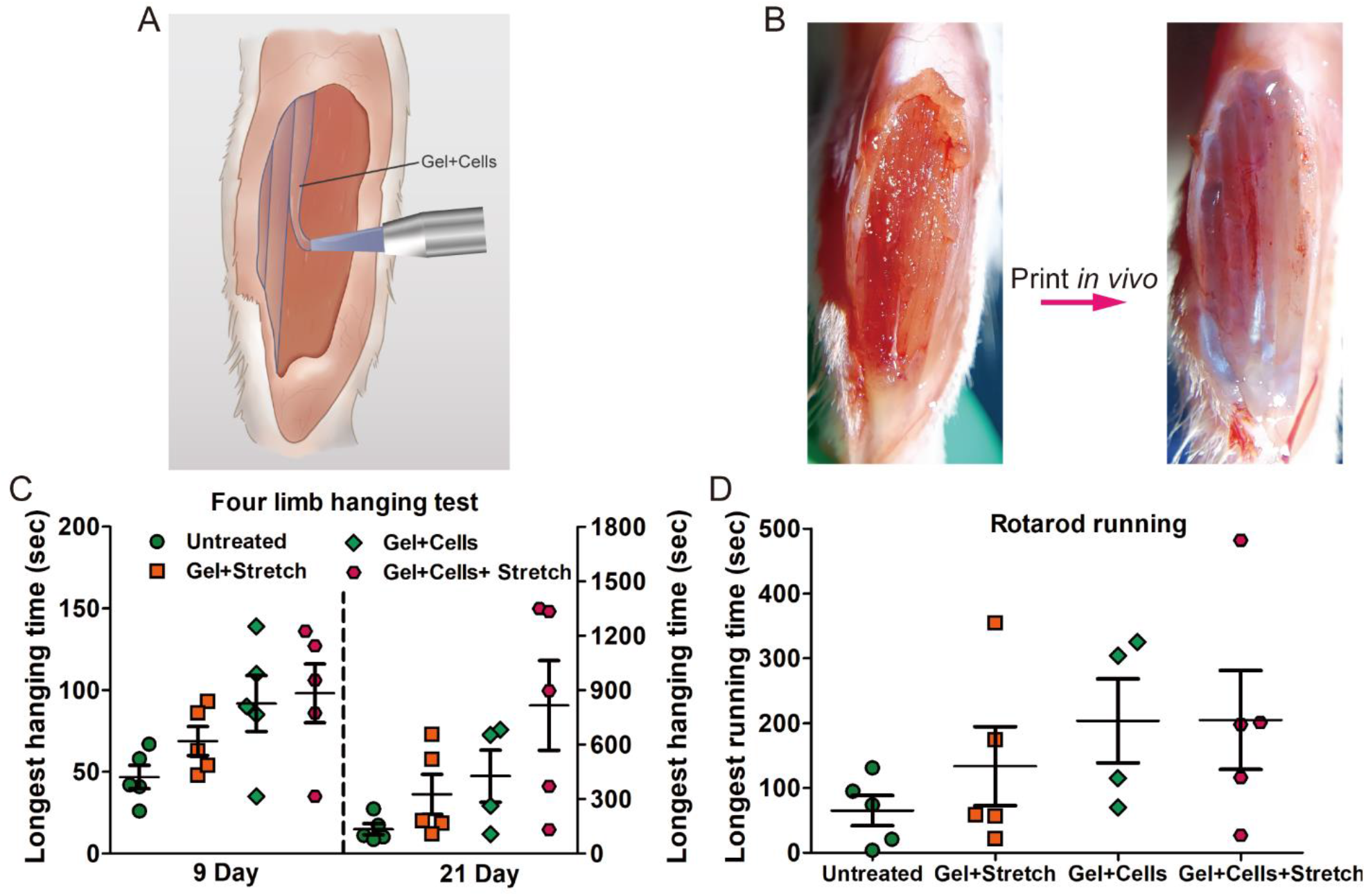
Print engineering myofibers in vivo to repair the VML damage. Schematic (**A**) and images (**B**) demonstrate the VML defects and myomimetic aligned gelatin filaments were printed onto wounds directly. Functional performance of mice assessed by limb hanging test (**C**) and rotarod running test (**D**).

**Fig. 6.**
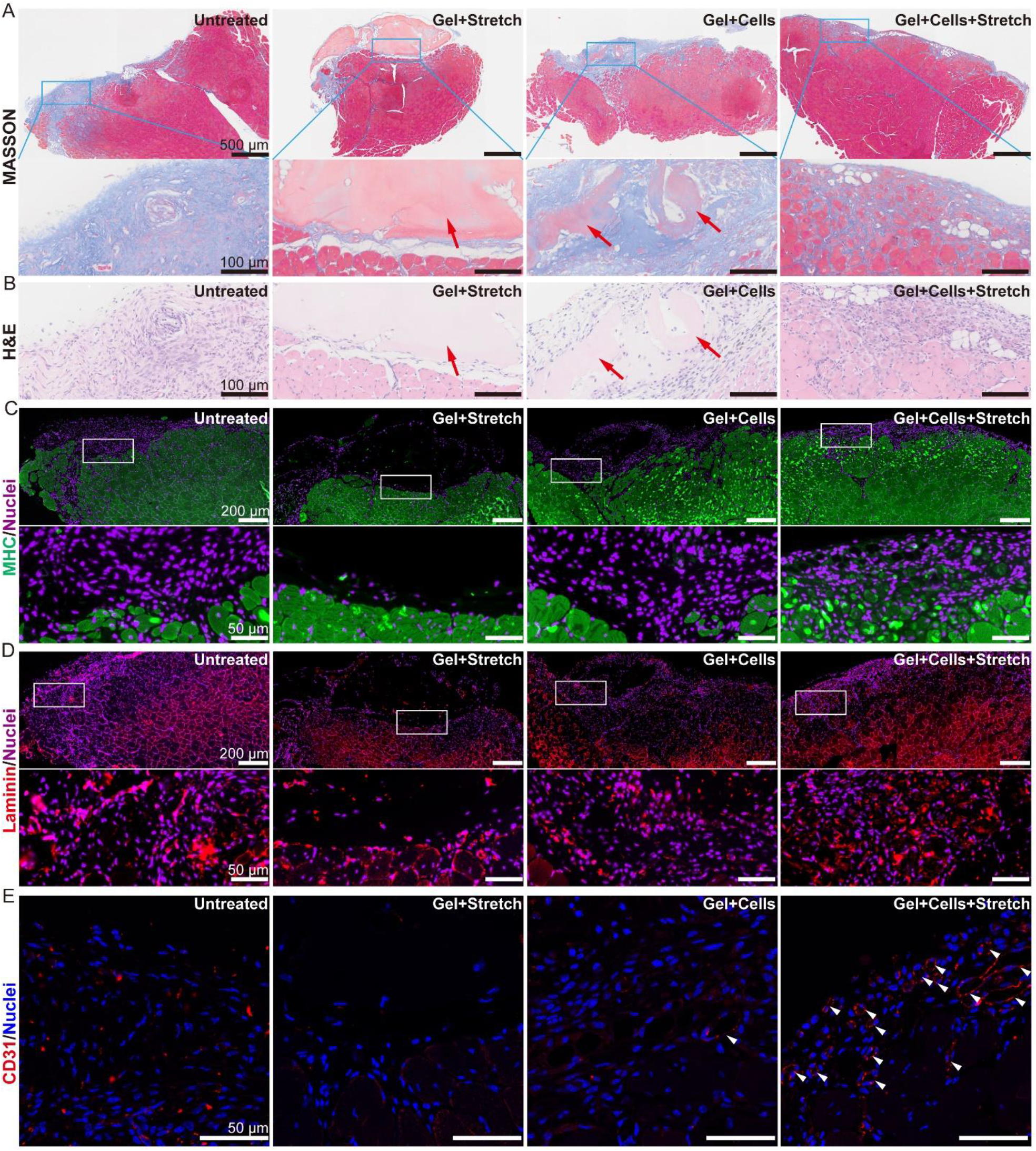
Tissue sections of repaired muscle. Masson’s Trichrome staining (**A**) and H&E staining (**B**) demonstrate much more regenerated (centrally-nucleated) myofibers and lower fibrosis within the defects in the experimental group (Gel+Cells+Stretch). Arrowheads denote residual gelatin. MHC (green) (**C**), Laminin (red) (**D**), and CD31 (red) (**E**) staining of VML defects treated with different methods at 3 weeks. Circular laminin structure and high densities of MHC+ myofibers and vessels (arrows) are present. Error bars: means ±SEM.

## Discussion and Conclusions

3D bioprinting is a research topic still in its early stages. Although the technique allows multiple biomaterials and different types of cells to be printed into constructs with desirable geometry and size, achieving the functionalities similar to those of the native tissues is a major challenge. Even for bioprinted tissues that have progressed to the preclinical stage, such as articular cartilage, reproducing their cell arrangements and thereby desired properties of the native tissues is yet to be accomplished (20). There is currently no reliable method that can consistently generate cell constructs which can reproduce the cell arrangements as well as the functionalities of native tissues. In this study, we have demonstrated a method to control cellular orientation in 3D bio-printed filaments. By integrating stretching into the extrusion bioprinting frame, we are able to create a mechanical environment which, in turn, would regulate the subsequent cellular alignment and differentiation in the generated tissues. This mechano-modulation phenomenon is broad based and was observed in several cell types in our experiments, suggesting an underlying common mechanism. We have investigated the effects of a few other mechanical stimuli on the cell responses such as shear stress and contact surface geometric patterns, and concluded that they did not influence the cell alignment. However, the underlying mechanism of how cells sense and response the static tensile stress is not very clear, and is currently being studied by our research group. This method offers a promising avenue for producing artificial tissues in vitro that possess the cellular arrangements and functionalities of native tissues.

If the fact that the mechanical microenvironment regulates cell orientation or even cell division is expected, the tensile stress induced myogenic fusion, differentiation and maturation of myoblasts without differentiation medium is surprising. These myogenic processes are generally believed to be triggered by biochemical cues. Our results show that there seems to be an equivalence between the biochemical and biophysical (mechanical) driving forces during myotube formation. More studies are needed to fully understand this “equivalence”. Our method lays out a foundation how the influence of mechanical stimuli on a number of biological processes of cells, such as cellular division (13, 14), differentiation (21), orientation and morphology (22), etc. can be investigated.

Our results on tensile stress-induced cellular orientation and division also provide clues for understanding tissue development and adult physiology. For instance, it is known that surgical incisions along the natural lines of skin tension would result in cosmetically pleasing surgical results (23). Although skin cleavage lines or skin tension lines have been used as guidance for elective surgical incisions over 100 years, the underlying cellular mechanisms are not understood (24). Our results show that cells, including human skin fibroblast (HFF-1), prefer aligning and dividing along the direction of the maximum tensile stress, providing a plausible explanation for the cosmetically smoother healing of surgical incisions. Additionally, in every type of investigated cells, we observed significant elongation. As we know, morphological changes are typically associated with changes of biochemical functions, for example, cell shape is associated with macrophage phenotype polarization and elongation of macrophage leads polarization toward prohealing (M2) phenotype (25). Therefore, the relationship between morphological changes and cellular biochemical functions in our system warrants further study.

Perhaps the most clinically promising result of the current study is the ability of the stretch-bioprinted filaments to repair damaged tissues, in particular, injured muscle in vivo. Such results not only suggest the essential roles mechanical stresses play in cell and tissue growth, but also point to the possibility of a full range of tissue engineering technologies in tissue regeneration, surgery, and wound repair applications.

## Materials and Methods

### Materials

Gelatin (from bovine skin), Pancreatin (form porcine pancreas), trypsin, 5-bromo-2-deoxyuridine (BrdU) were acquired from Sigma Aldrich. Sulfosuccinimidyl 6-(4’-azido-2’-nitrophenylamino) hexanoate (Sulfo-SANPAH), Hanks’ Balanced Salt Solution (HBSS 1X), fetal bovine serum (FBS), Dulbecco’s Modified Eagle Medium/Nutrient Mixture F-12 (DMEM/F-12) were purchased from Life Technologies. Phalloidin-YF555 was purchased from US Everbright Inc (Suzhou). Phalloidin-iFluor 488, anti-Vimentin antibody (ab92547), anti-alpha smooth muscle actin antibody (ab124964), anti-fast myosin skeletal heavy chain antibody(ab51263), anti-laminin antibody (ab11575), anti-CD31 antibody (ab28364), secondary antibody (ab150078, Alexa Fluor 555) were purchased from Abcam. Collagenase II was acquired from Worthington. Collagen I was acquired from Corning. Gelatin Methacryloyl (GelMA), lithium phenyl-2, 4, 6-trimethylbenzoylphosphinate (LAP) were acquired from Suzhou Intelligent Manufacturing Research Institute (Suzhou, China). Microbial transglutaminase (mTG) were acquired from Cool Chemical Technology (Beijing) Co., Ltd. Cytochalasin B, fibrinogen and 4’, 6-diamidino-2-phenylindole (DAPI) were bought from Solarbio Life Science (Beijing). Human skin fibroblast (HFF-1) was bought from Cobioer Biosciences (Nanjing) co., ltd. Thrombin and red blood cell lysis buffer was bought from Beyotime Biotechnology (Shanghai). Polyacrylamide gel (SurePAGE™, 4-20%) was bought from Genscript Biosciences (Nanjing) co., ltd. Blebbistatin was bought from Macklin Biochemical (Shanghai) Co., Ltd.

### Methods

#### Extraction of Neonatal Rat Cardiomyocytes and Cardiac Fibroblasts

All animal procedures were done in accordance with the guidelines of the Animal Use and Care Committee of Zhejiang University (China). Ventricles were removed from 2-day-old Sprague Dawley rat pups (Zhejiang Academy of Medical Sciences) and the tissue was manually minced into pieces sized around 0.5-1 mm3 and placed in a Hank’s balanced salt solution containing 0.07% trypsin, 0.02% pancreatin and 0.05% collagenase for 8 min of enzymatic digestion at 37 °C for 10 times, followed by digestion termination with DMEM/F-12 medium solution (20% FBS). The turbid liquid was passed through a 70 μm cell strainer. Cardiomyocytes, cardiac fibroblasts and red blood cells were further isolated by centrifuging at 800 rpm for 5 min. Subsequently, the cells were dispersed into DMEM/F-12 solution which contained 20% FBS and 0.1 mmol BrdU, and precultured in cell culture bottle to separate cardiac fibroblasts from cardiomyocytes. After nearly 90 min, the unattached primary cardiomyocytes and red blood cells were extracted by centrifuging at 800 rpm for 5 min, while cardiac fibroblasts were retained in the culture bottle. Then, red blood cells were removed with red blood cell lysis buffer. Cardiomyocytes were harvested for subsequent experiments and cardiac fibroblasts were cultured in DMEM/F-12 medium which containing 20% FBS and 1% penicillin-streptomycin in constant temperature (37 °C) and humidity incubator with 5% CO2.

#### Extraction of Rat Airway Smooth Muscle Cells

After anesthesia, two male Sprague-Dawley rats (about 50 g) were sacrificed by cervical dislocation, soaked in 75% ethanol and sterilized for 2 min. Then the trachea and lung tissues were quickly separated and placed in HBSS buffer containing 1% penicillinstreptomycin. Lung tissues and remaining connective tissues were removed. After soaking the trachea with dispase II(10 mg/mL)for 45 min, the inner and outer membrane of the trachea was stripped off. Then the trachea was cut into 1 mm3 pieces and then incubated in digestive solution (5 ml HBSS containing 31 mg collagenase I, 2 mg papain, 10 mg bovine serum albumin and 0.7 mg dithiothreitol) at 37°C for 25 min, serum-containing medium was added to terminate the digestion. The turbid liquid was passed through a 70 μm cell strainer, centrifuged at 1000 rpm for 5 minutes. The cells were resuspended in the culture medium and seeded in the T25 culture flask. After grown to 80% confluency, the cells were digested with 0.25% trypsin protease-EDTA and centrifuged for 5 min. After centrifugation, cell pellet was resuspended and transferred to a new T25 culture flask. After 30-60 min, the unattached airway smooth muscle cells were carefully aspirated and transferred to a new culture flask. Then purer smooth muscle cells can be obtained, which can be used for about 8 generations.

#### Extraction of Rat Dorsal Root Ganglion (DRG) Neurons

5 Sprague-Dawley rats (5-to 8-day-old) were sacrificed by decapitation, and the skin overlying the spinal cord was cut away. Detach the spine from the body of the pups and then cut along the long axis of the spine to open the spine completely and extract the ganglion from the bone pocket. Then, transfer the ganglia to 5mL of ice-cold HBSS buffer containing 1mL penicillin/streptomycin. Let the tissues settle to the bottom of the tube and gently remove the HBSS. Next, add 10 ml of dissociation medium (9 mL of 0.05% Trypsin-EDTA and 1 mL of 500 U/mL collagenase A) into the tube. Incubate it at 37 °C for 90 min while shaking the tube every 10 min. After the dissociation, centrifuge the ganglia at 900 g for 5 min and remove the supernatant. The pellet is further re-suspended in 10mL of standard growth media and centrifuged at 900 g for 5 min. After washing with the medium, the cells are re-suspended in standard growth media containing the Neurobasal-A medium, B-27 supplement (2%), penicillin/streptomycin (0.5%), nerve growth factor (5 μg/mL), fetal bovine serum (1%) and glutamax (1%) and cultured in constant temperature (37 °C) and humidity incubator with 5% CO2.

#### Manufacturing 3D Patterned Constructs with Control of Cellular Orientation

Cells aligned in the interior of gel: 3 million cells were resuspended in 900μL 6.6% gelatin/ (DMEM/F12) and then mixed with 100 μL 50mg/mL mTG/ HBSS. Then the cell suspension was transferred into the teflon tube (600 μm internal diameter) of our device with syringe immediately, followed by setting the device at room temperature for gelation. After 45 minutes, gel formed, and then gel-cell mixture was forced out of teflon tube by pushing the syringe slowly. Since gelatin could adhere to polystyrene substrate, one end of the gelatin filament could be fixed onto the plate when it encountered with the substrate. Next, we could stretch the gelatin and print desired constructs merely by moving the microextrusion head. These generated constructs were cultured in corresponding cell culture medium containing 20% FBS and 1% penicillin-streptomycin in constant temperature (37 °C) and humidity incubator with 5% CO2

De-stretch: First, gelatin filaments were printed onto PDMS substrate under “dry” condition. Then, within 20 minutes, culture medium was added. Once encountered with medium, gelatin filaments detach themselves from the PDMS substrate.

Cells aligned on the surface of gel: The procedure of constructing the acellular constructs is the same as above. After these patterned constructs formed, cells were seeded evenly on the surface of them.

GelMA gel: Most of the process for fabricating patterned GelMA gel is the same as that for manufacturing patterned gelatin gel, except that 4% GelMA (containing 0.5% (w/v) LAP) took 15 seconds of UV light (3 W, 405 nm) to gelation before extrusion rather than enzyme-catalyzed gelation.

#### VML Defects Model

Animal and surgical procedures were approved by Animal Use and Care Committee of Zhejiang University (China). Bilateral VML defects in female NOD-scid IL2Rgnull (NSG) immunodeficient mice (8 weeks, GemPharmatech Co., Ltd) were used for all experiments. Mice were anesthetized with pentobarbital sodium, then the tibialis anterior muscle was exposed, and approximately 30% (about 13 mg) of the muscle was removed. Cells loaded gelatin or pure gelatin were printed onto the wounds. For another group, cells loaded gelatin was dropt onto the wounds. The structures were fixed using fibrin glue and the skin incision site was sutured (6-0 suture). Three weeks after surgery, the entire tibialis anterior muscles were harvested and fixed using 10% formalin solution, which followed by paraffin embedding for histological evaluation (hematoxylin and eosin staining, Masson’s Trichrome staining and immunofluorescence staining).

#### Cellular Immunofluorescence Staining

Samples were fixed in PBS with 4% v/v paraformaldehyde for 20 min and immersed in 0.1% v/v Triton-X100/PBS solution for 15 min to permeabilize cytomembrane. Then, samples were incubated in blocking solution (2%BSA, 4% goat serum) for 30 min. Subsequently, staining with primary antibody (1:200 dilution) was conducted for 12 hours at 4 °C, and an additional staining with secondary antibody (1:200 dilution), DAPI (5 mg/mL), and phalloidin-iFluor 488 (1:1000 dilution) was conducted for 2 hours at room temperature. Between each step, the samples were washed by PBS three times. All antibody incubations were performed in blocking solution.

#### Characterizing the Surface and Section of the Stretched Gelatin Filaments with SEM

Surface: Stretched gelatin filaments were fabricated onto PDMS and freeze by using liquid nitrogen immediately. Then, these filaments were lyophilized in a vacuum-freeze dryer before they were observed by scanning electron microscopy.

Section: Stretched gelatin filaments were fabricated onto the frozen cryo-embedding media (Tissue-Tek, O.C.T., Sakura, The Netherlands) directly. Subsequently, these filaments were coated with cryo-embedding media and frozen at −20°C and were sliced into 10-μm-thick sections by using a cryostat freezing microtome. Then, these sections were transferred onto PDMS and were freeze in liquid nitrogen immediately. After lyophilized, these sections were observed by scanning electron microscopy.

#### Pharmacological Cytoskeleton Inhibitors

C2C12 and HFF-1 cells suspension were mixed with pharmacological cytoskeleton inhibitors, and then were seeded onto the surface of stretched gelatin. Concentrations of 10 μM blebbistatin and 5 μM Cytochalasin B were used and cells were incubated with them respectively for 1 day at standard culture conditions before microscopic observation.

#### Characterization

SEM images were obtained by a scanning electron microscope (Hitachi SU-8010). Confocal microscopy images were taken by a laser-scanning microscope (Leica, TCS SP8). Images of paraffin sections were obtained with Olympus VS120. The Cell orientation distribution and cell length were quantified by using ImageJ.

## Acknowledgments

We thank Xiaoqiu Shu, Tingting Ye and Qunchen Yuan for helpful discussions. We also thank for the technical support by the core facilities, Zhejiang University School of Medicine.

## Author Contributions

Chuanjiang He, Ping. Wang, and K. Jimmy. Hsia designed the experiments. Chuanjiang He performed the experiments and analyzed the data with help from Mengxue Liu, Deming Jiang, Tao Liang, Pan Wu and

Chunmao Han. Chuanjiang He, Liquan Huang, K. Jimmy. Hsia and Ping Wang wrote the manuscript with input from Mengxue Liu and Chunlian Qin. Ping Wang acquired the financial support and led the project;

## Data and materials availability

All data is available in the main text or the supplementary materials. The raw image data and the analyze data generated in this study are available from the corresponding author on reasonable request.

## Supplementary Materials

**Fig. S1.**
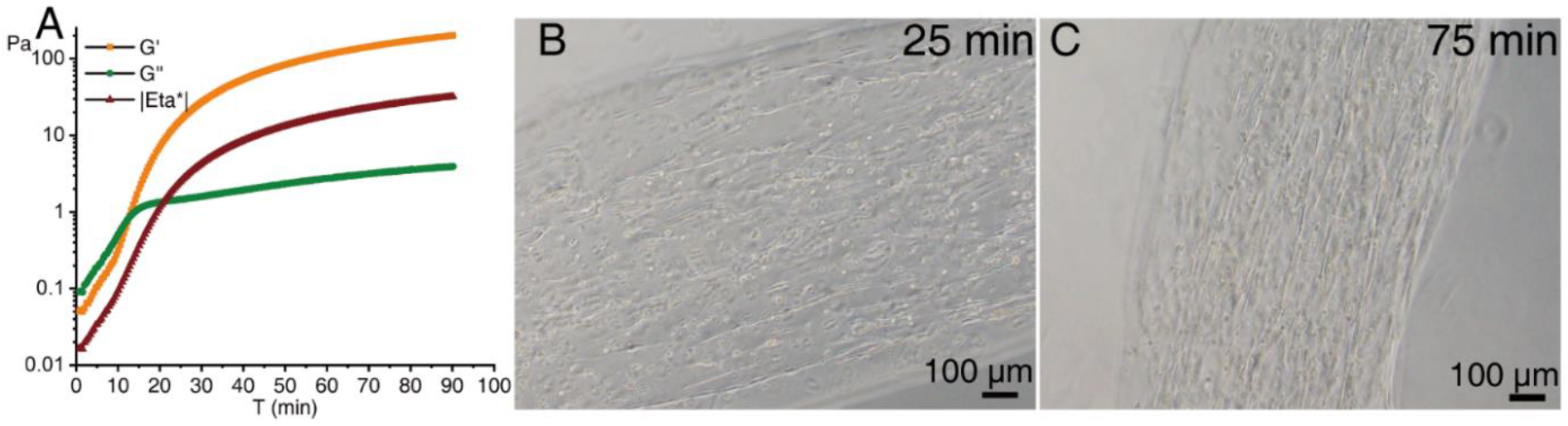
Cells aligned in the stretched gelatin which subjected with different gelation time. (A) Evolution of the storage modulus G’, loss modulus G” and complex viscosity (|Eta *|) during gelation of 6% gelatin at 25 °C. Before reaching the gel point, G” is higher than G’, indicating the liquid state of gelatin. Beyond the gel point, G’ is higher than G”, which is a proof of solidification. Cardiomycytes aligned in the interior of stretched gelatin gel after 25 min (B) and 75 min (C) gelation.

**Fig. S2.**
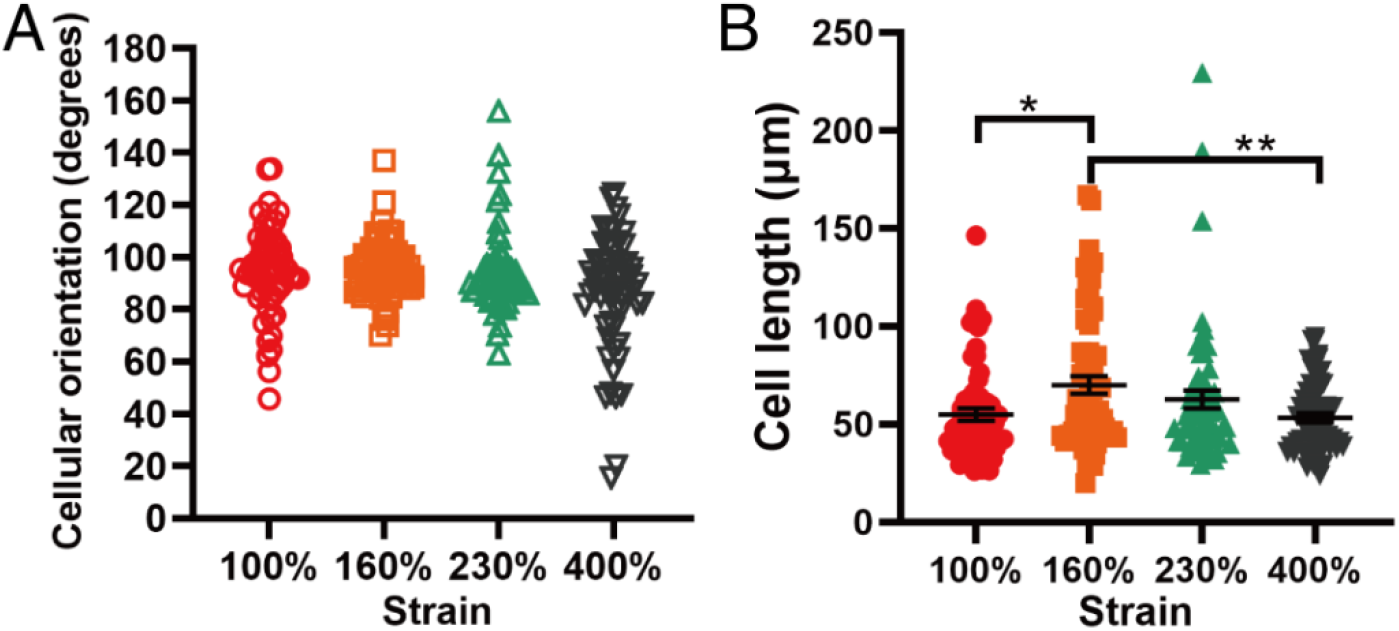
The cellular orientation distribution (A) and cell length (B) of cardiomyocytes in the interior of the stretched gelatin which subjected with different strain. The 90° angle represents the direction of stretching. n=60, Error bars: means ± SEM, one-way ANOVA with Tukey’s multiple comparisons test, *p = 0.024, **p = 0.0094

**Fig. S3.**
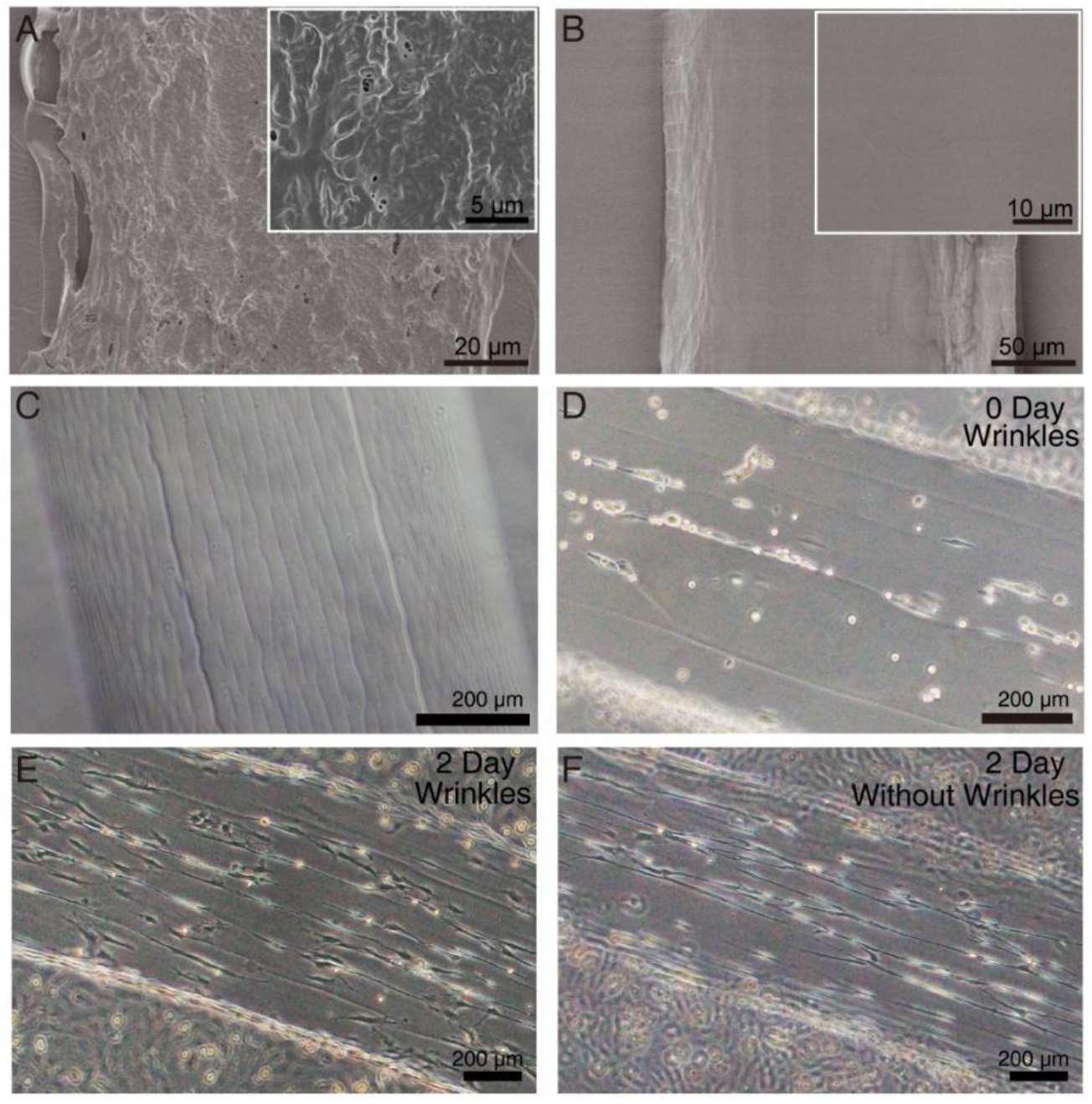
Role of anisotropic topography in inducing cellular orientation. SEM images of the section (A) and surface (B) of lyophilized stretched gelatin. Upper right: magnified images. (C) Wrinkles generated by air-drying process. (D) Cardiac fibroblasts prefer to fall into the wrinkles on the stretched gelatin. Cardiac fibroblasts seeded on the surface of stretched gelatin with (D) and without (E) wrinkles for 2 days.

**Fig. S4.**
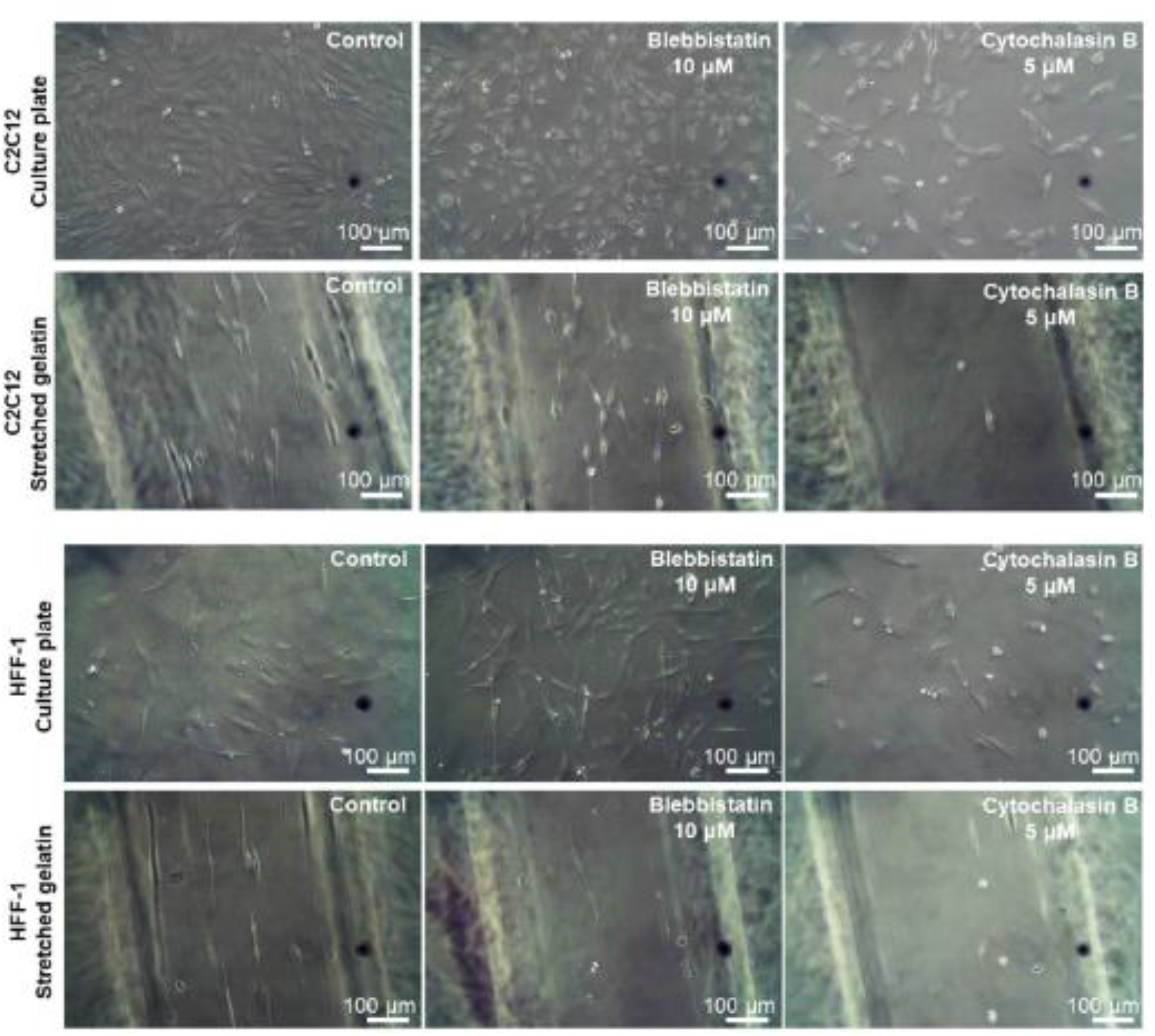
Cellular adhesion and alignment were inhibited by pharmacological cytoskeleton inhibitors, blebbistatin and cytochalasin B.

**Fig. S5.**
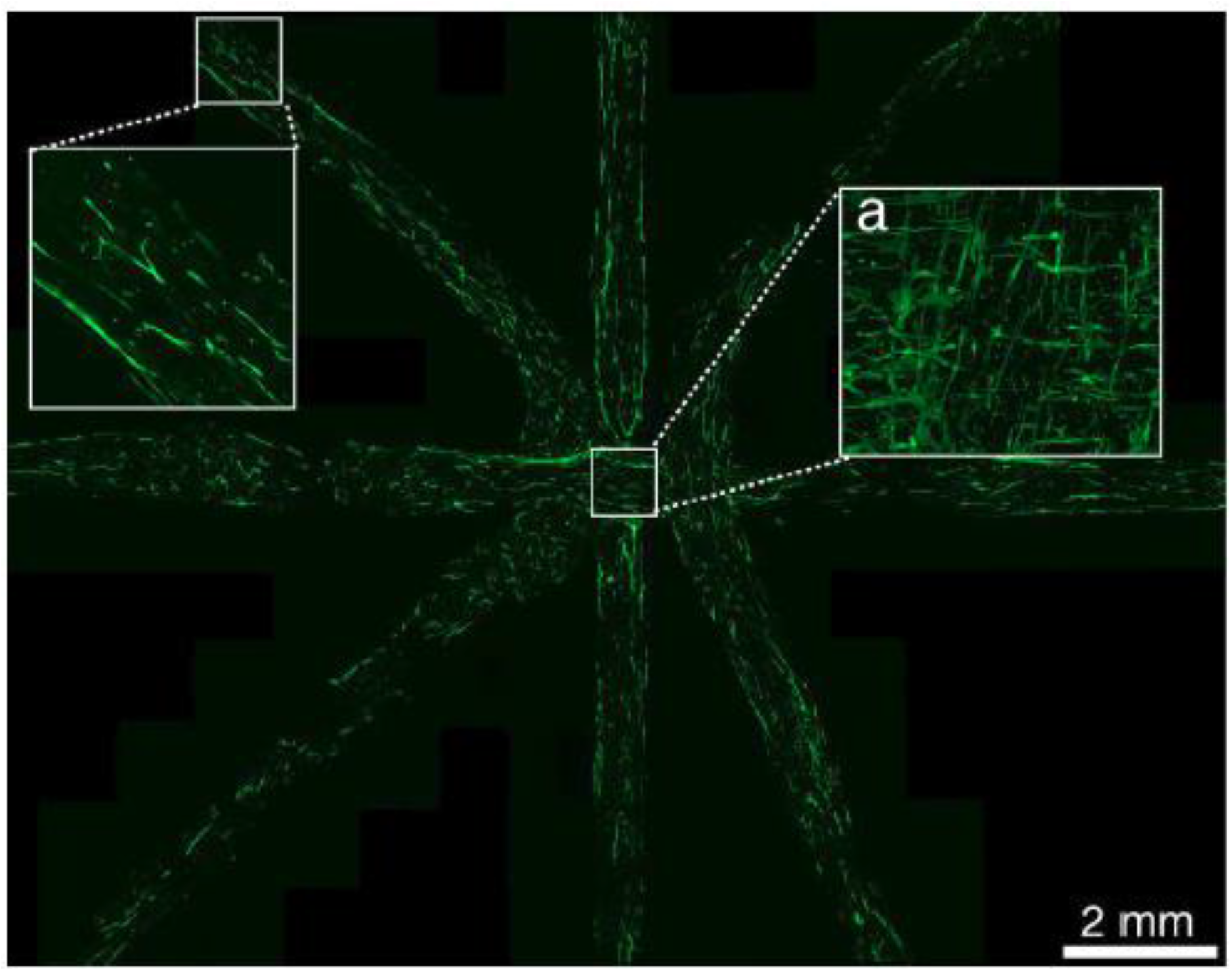
Confocal section image of cardiomyocytes aligning in the 3D patterned gelatin. (a). Maximum z-projection image of the box region. Green: phalloidin- iFluor 488.

**Fig. S6.**
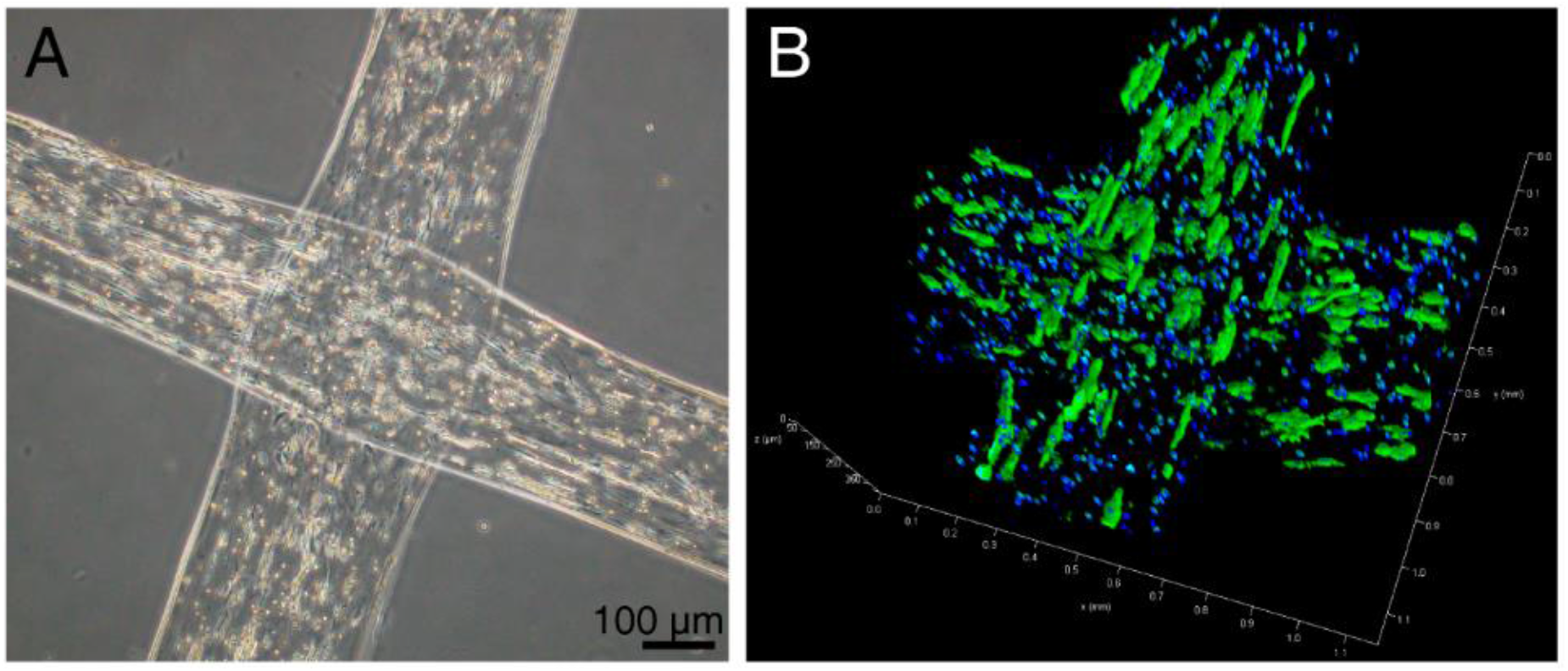
Vertically stacked constructs printed with cells-loaded gelatin. (A) Image of cardiomyocytes arranged in the interior of the crisscross construct printed with 6% gelatin. (B) Confocal fluorescence 3D image of cardiomyocytes arranged in the interior of the crisscross construct fabricated with 8% gelatin. Green: phalloidin- iFluor 488; Blue: DAPI.

**Fig. S7.**
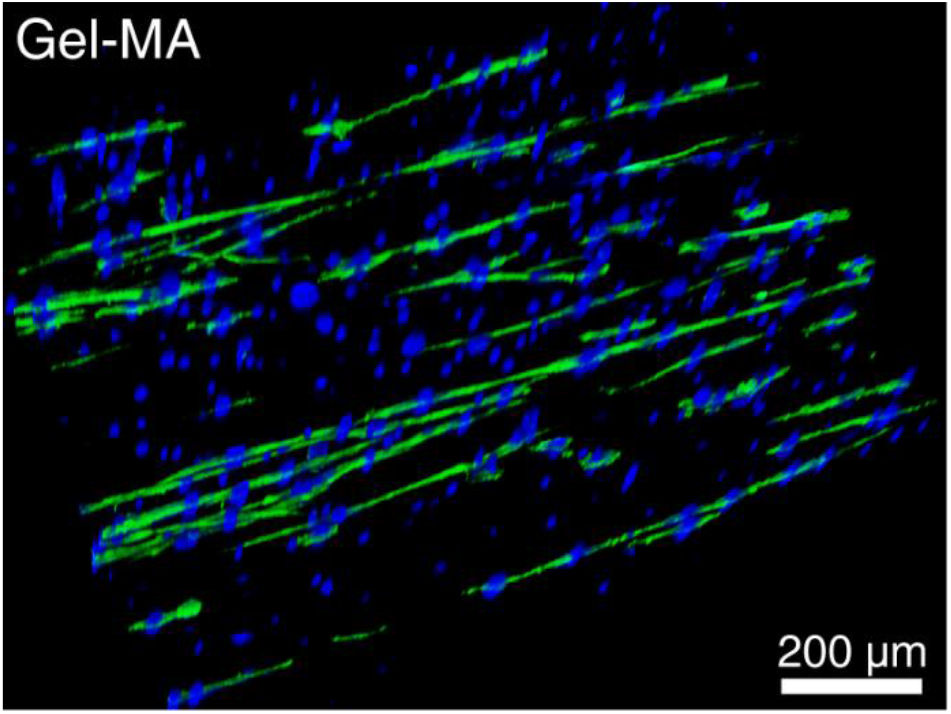
Confocal fluorescence 3D image of cardiac fibroblasts aligning in printed GelMA constructs. Green: phalloidin- iFluor 488; Blue: DAPI.

**Fig. S8.**
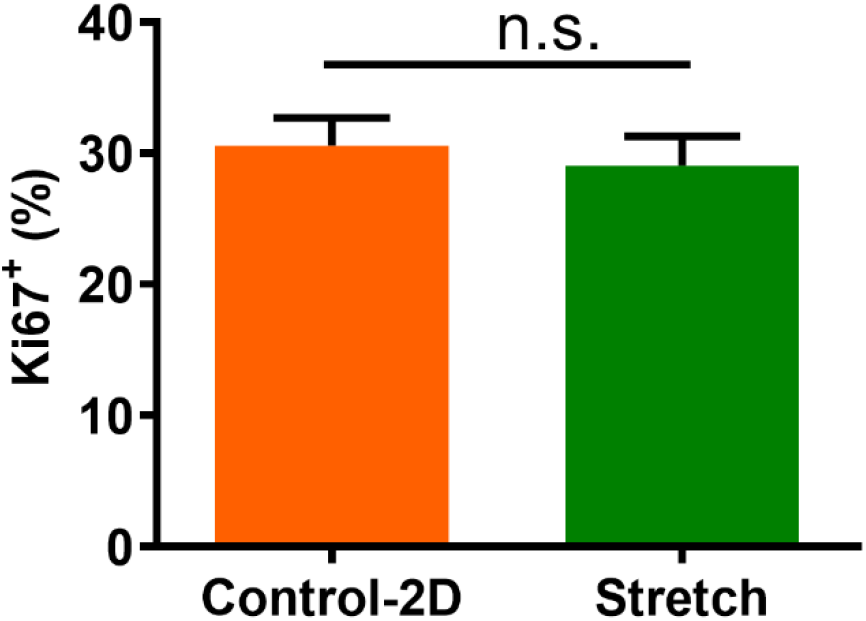
Proliferation (Ki67^+^) rate of C2C12 cells in stretched gelatin constructs and on the substrate of culture plates. Error bars: means ± SEM, unequal variance Student’s t-test, p= 0.6397.

**Table S1.** Mouse body weight and tibialis anterior muscle weight before and after the experiment. Values are means ± SEM.

| Treatment Group | Initial Body<br>Weight(g) | Body Weight at<br>the End (g) | Resected Muscle<br>Weight (mg) | Harvested<br>Muscle Weight<br>(mg) |
| --- | --- | --- | --- | --- |
| Untreated | 21.44±0.47 | 20.61±1.00 | 14.78±0.48 | 27.92±1.20 |
| Gel+Stretch | 20.1±0.47 | 19.06±0.38 | 12.84±0.81 | 29.78±1.92 |
| Gel+Cells | 21.28±0.38 | 19.91±0.77 | 13.71±0.49 | 32.825±3.12 |
| Gel+Cells+Stretch | 21.94±0.29 | 20.57±0.58 | 13.1±0.43 | 37.44±1.09 |

### Movies S1 to S3

Movie S1. Cardiomyocytes aligned in the interior or on the surface of stretched gelatin

Movie S2. Engineering myofibers with myomimetic hierarchical structure

Movie S3. The process of printing and aligning engineering myofibers directly onto the injured muscle

